# ERK5 is required for neutrophil-mediated ROS release and a key contributor in epidermolysis bullosa acquisita

**DOI:** 10.1101/2025.11.12.688045

**Authors:** Sören Dräger, Mareile Schlotfeldt, Leonie Voss, Sophia Johannisson, Michael Radziewitz, Colin Osterloh, Natalie Gross, Mayumi Kamaguchi, Ann-Kathrin Schneider, Gestur Vidarsson, Remco Visser, Frank Petersen, Xinhua Yu, Andreas Recke, Nancy Ernst, Anja Lux, Katja Bieber

## Abstract

Neutrophils are key effector cells in antibody-mediated autoimmune diseases, contributing to inflammation via the release of reactive oxygen species (ROS). Extracellular signal-regulated kinase 5 (ERK5), a member of the MAPK family, is expressed in neutrophils but its role in autoimmune disease pathogenesis remains unclear. We investigated the functional relevance of ERK5 in antibody-mediated autoimmune diseases by comparing neutrophil-dependent (epidermolysis bullosa acquisita, EBA; serum transfer arthritis, STA) and neutrophil-independent (immune thrombocytopenia, ITP) murine models using the small-molecule ERK5 inhibitor XMD8-92. *In vitro*, pharmacological ERK5 inhibition specifically reduced neutrophil ROS release and CD62L shedding without affecting adhesion, chemotaxis, or CD18 expression. No major effects on viability were observed. *In vivo*, ERK5 inhibition with XMD8-92 significantly ameliorated antibody transfer-induced EBA, supporting a critical role of neutrophil-derived ROS in disease pathogenesis. In STA and ITP, ERK5 inhibition did not affect clinical outcomes. Together, these findings highlight ERK5 as a regulator of neutrophil ROS release and a potential therapeutic target in ROS-driven autoimmune diseases such as EBA.

## Introduction

Neutrophils are the most abundant granulocytes and key effector cells of the innate immune system (Zhang et al. 2024). They are recruited from the circulation to the site of inflammation through a multi-step process (Springer 1994), including selectins, chemoattractants and integrins and - in the context of autoimmune diseases – are activated by deposited antibodies/immune complexes (ICs) via Fc gamma receptors (FcγR) (Ludwig et al. 2017). In the context of antibody-mediated diseases, e.g. autoimmune blistering disorders such as pemphigoid diseases, human neutrophils are activated via FcγRIIA and FcγRIIIB (Ludwig et al. 2017). Correspondingly in mice, FcγRIV is critical for pathogenesis (Kasperkiewicz et al. 2012), whereas FcγRIIB is inhibitory (Karsten et al. 2012). FcγR activation by IgG leads to internal signal transduction via spleen tyrosine kinase (SYK), transforming growth factor-β-activated kinase 1 (TAK1) and the mitogen activated protein kinase (MAPK)/mitogen-activated protein kinase kinase (MEK)/extracellular signal-regulated kinases (ERK) signaling cascade (Zhang et al. 2024). Consequently, inhibiting these pathways can alleviate disease, which has been shown for example for SYK (Samavedam et al. 2018) or PI3K (Ghorbanalipoor et al. 2022; Koga et al. 2018; Zillikens et al. 2021) in murine models of epidermolysis bullosa acquisita (EBA). Notably, a recent high-throughput screening of a drug library identified several repurposable drugs targeting polymorphonuclear leukocytes (PMNs) (Ghorbanalipoor et al. 2024). Since neutrophils contribute to the inflammation with their effector functions, such as the release of reactive oxygen species (ROS) (Chiriac et al. 2007a; Shimanovich et al. 2004) and proteases, such as neutrophil elastase (Liu et al. 2000) or granzyme B (Emtenani et al. 2024), this opens possibilities for further treatment exploration.

As ERK5 is expressed in neutrophils (Hii et al. 2004) and upregulated upon activation (Zillikens et al. 2021), it represents a promising target for therapeutic intervention in diseases primarily driven by neutrophils. ERK5, encoded by the *MAPK7* gene and also known as big MAP-kinase 1 (BMK1), is a member of the MAPK family, which are serine/threonine kinases centrally involved in a range of cellular processes including proliferation, differentiation, migration, responses to stress, cell survival and regulation of inflammation (Cargnello and Roux 2011). Whereas the classical MAPK pathways including ERK1/2, p38, and JNK pathways are already extensively characterized in neutrophil immune responses and autoimmune diseases (Müller et al. 2016; Vorobjeva 2023; Zarrin et al. 2021a), the function of ERK5 in this context is largely unknown. However, in other cell types such as endothelial cells and monocytes, ERK5 is associated with e.g. oxidative stress (Tusa et al. 2023), inflammation (Wilhelmsen et al. 2015) and survival signaling (Gírio et al. 2007). Since these processes also play a role in activated neutrophils (Aroca-Crevillén et al. 2024), we hypothesize that ERK5 is involved in immune complex-induced kinase signaling in neutrophils. It acts downstream of MAP kinase kinase kinase (MEKK2/3) and MAPK2 MEK5, and is engaged by a variety of stimuli, e.g. cytokines, hormones, mitogens and stress-factors like shear stress and high osmolarity (Le 2023). In its closed inactive conformation, ERK5 is complexed with heat shock protein 90 (HSP90) and the co-chaperone cell division cycle 37 (CDC37), exposing a nuclear export signal (Erazo et al. 2013; Kondoh et al. 2006). Upon phosphorylation by MEK5, cyclin dependent kinase (CDK) 5, CDK1 or ERK1, ERK5 auto-phosphorylates several residues within its C-terminal extension, resulting in conformational changes, dissociation from HSP90 and exposure of the formerly hidden nuclear localization signal (Gomez et al. 2016). After transport into the nucleus, ERK5 mediates phosphorylation of MEF2 transcription factors and also acts as a transcription factor itself (Kasler et al. 2000; Morimoto et al. 2007). The pathophysiological role of ERK5 in disease has been extensively reviewed elsewhere (Le 2023; Paudel et al. 2021). ERK5 has primarily been investigated in cancer, such as hematological malignancies and melanoma, where ERK5 inhibition induced senescence of melanoma cells (Tubita et al. 2022). However, its role in autoimmune disorders is underexplored, with only few published studies on this topic. For example, in psoriasiform dermatitis, treatment with the ERK5 inhibitor AX-15836 reduced disease severity and inhibited neutrophil extracellular trap (NET) formation (Ding et al. 2022). ERK5 is also required for B cell activating factor (BAFF)-induced B cell survival (Jacque et al. 2015), hence, inhibition of ERK5 might be beneficial in antibody-mediated autoimmune diseases.

Based on the aforementioned data and the prominent role of neutrophils in the effector phase of antibody mediated diseases, we designed this study to clarify the role of ERK5 in antibody-mediated autoimmune diseases by contrasting neutrophil-dependent and -independent conditions. To highlight pathogenic differences and to assess the impact of ERK5, we selected three murine models representing prototypical antibody-mediated autoimmune diseases. EBA and serum transfer arthritis (STA) (as a model for rheumatoid arthritis (RA)) were chosen as examples of neutrophil-driven diseases. Due to distinct involved autoantigens, EBA is characterized by dermal-epidermal separation leading to skin blistering (Tigges et al. 2025) and STA by swelling of joints accompanied by bone and cartilage destruction(Christensen et al. 2016). The pathogenesis of both diseases has been shown to be highly neutrophil-dependent (Chiriac et al. 2007a; Shimanovich et al. 2004; Wipke and Allen 2001) but in contrast to EBA, an additional role for macrophages, mast cells, and NKT cells has been shown in STA (Christensen et al. 2016). To further delineate a possible neutrophil-specific effect, the impact of ERK5 inhibition in immune thrombocytopenia (ITP) was evaluated, a disease model considered neutrophil-independent (Biburger et al. 2011). Here, platelet-bound IgG autoantibodies induce immune cell signaling mainly through FcγRI and FcγRIV (Biburger et al. 2011), resulting in the subsequent phagocytosis and destruction of platelets in reticuloendothelial organs driven by tissue-resident macrophages (Biburger et al. 2011; Nimmerjahn et al. 2005; Norris et al. 2023; Semple 2010; Wöhner et al. 2024; Zufferey et al. 2017).

The use of murine disease models has been invaluable in advancing our understanding of the disease pathogenesis and in uncovering potential therapeutic strategies (Bieber et al. 2010; Dräger et al. 2017; Samavedam et al. 2018; Tigges et al. 2025; Zillikens et al. 2021). Aiming to uncover the role of ERK5 in IgG-driven autoimmunity, we herein investigate the impact of pharmacological ERK5 inhibition in antibody-mediated neutrophil-dependent (EBA and RA) and neutrophil-independent (ITP) autoimmune disease models.

## Results

### ERK5 inhibition selectively reduces ROS release from immune-complex–stimulated neutrophils

Neutrophil activation and recruitment to the inflamed tissue is a complex multistep process (Kolaczkowska and Kubes 2013). To assess the role of ERK5 in various neutrophil effector functions, we conducted several *in vitro* assays focusing on the following characteristics: ROS release, expression of the surface activation markers CD18 and CD62L, adhesion, and chemotaxis. Furthermore, potential toxic effects on neutrophils were investigated. In each assay, four different concentrations of the ERK5 inhibitor XMD8-92 (10 µM, 1 µM, 0.1 µM, and 0.01 µM) were used. While we could not show any differences in either the surface expression of CD18, adhesion, nor chemotaxis, ROS release was significantly reduced by the ERK5 inhibitor in a dose-dependent manner, with an IC_50_ of 1.469 µM. In addition, less CD62L shedding was observed at 10 µM and 0.01 µM. Of note, viability was slightly lower at 10 µM. However, since less than 1 % of the cells were affected, the biological significance is probably negligible, thus ruling out toxicity as main driver. (Figure 1 A-F).

**Figure 1.**
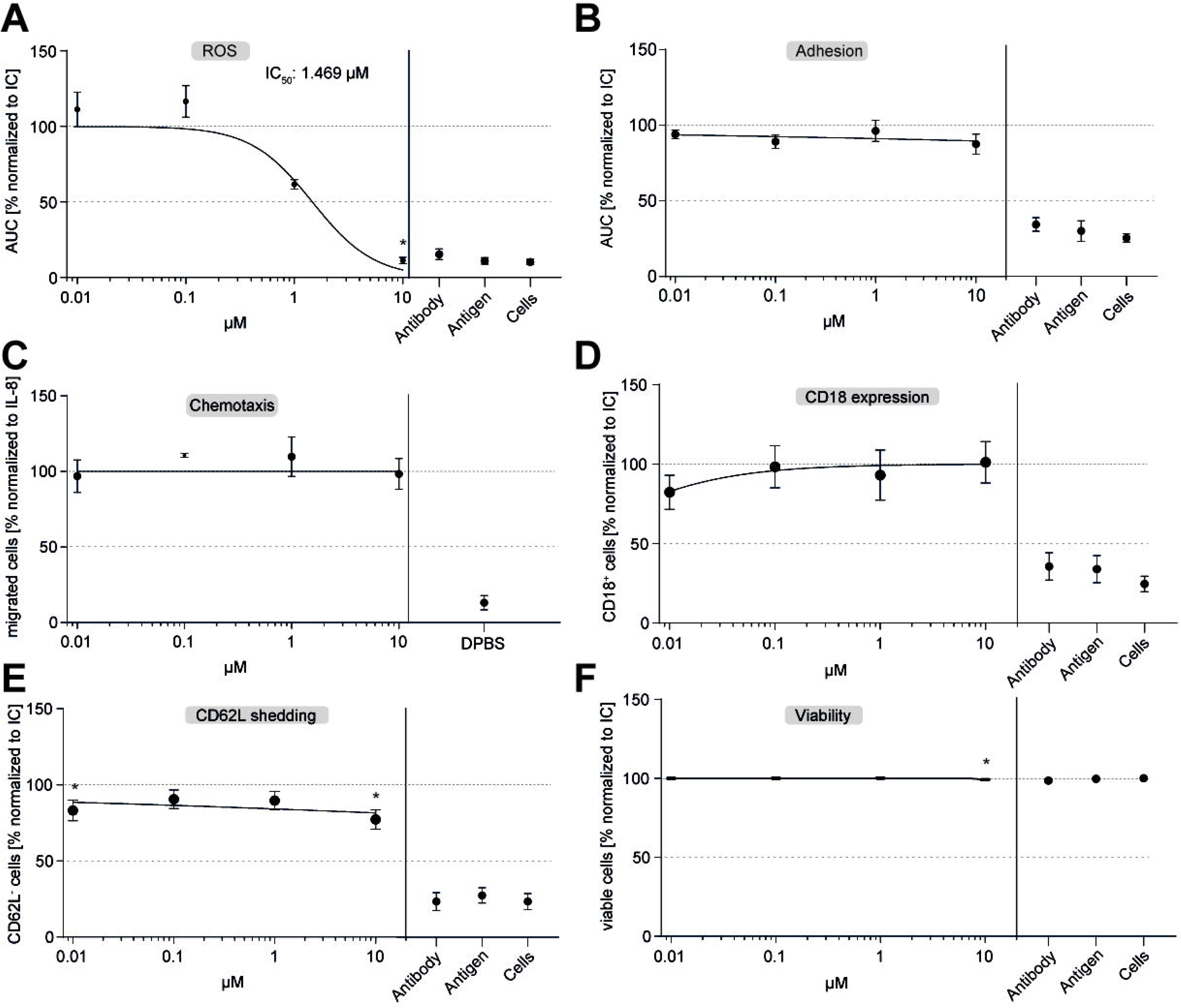
ERK5 inhibition selectively reduces ROS release and CD62L shedding from immune-complex (IC)–stimulated polymorphonuclear leukocytes (PMNs). Different cellular functions of neutrophil granulocytes were measured using increasing concentrations of the ERK5 inhibitor XMD8-92 (10 µM, 1 µM, 0.1 µM, 0.01 µM). **(A)** IC-induced ROS release was measured using a luminol-based assay, showing a dose-dependent reduction (IC□□ = 1.469 µM). **(B)** PMN adhesion upon IC-stimulation was investigated using impedance-based measurements, which revealed no significant changes. **(C)** Boyden chamber chemotaxis to IL-8 was measured using the myeloperoxidase assay. Chemotaxis was not altered by ERK5 inhibition. Surface expression of the cell activation markers, measured by flow cytometry of **(D)** CD18 and **(E)** CD62L shedding was reduced at 0.01 µM and 10 µM. **(F)** Cell viability, determined by staining for Annexin V and Zombie NIR, was slightly affected at 10 µM. All results were normalized to the respective controls without the ERK5 inhibitor. n = 5. Values are displayed as mean ± SEM. A nonlinear fit dose response curve was calculated. Kruskal-Wallis test with Dunn’s multiple comparisons test. *p ≤ 0.05.

### ERK5 is activated in lesional skin of EBA mice, and its inhibition by XMD8-92 alleviates disease development without affecting keratinocyte C5a release or antibody deposition

Kinases are crucially involved in the signaling of a variety of cellular responses in health and disease, including autoimmune diseases (Zarrin et al. 2021b). Therefore, we hypothesized that ERK5 is critically involved in the pathogenesis of EBA, which is a neutrophil-driven autoimmune skin disease (Koga et al. 2019). To investigate this, we measured the phosphorylated and thus active ERK5 (pERK5) in skin samples obtained from the immunization-induced EBA mouse model. pErk5 was subjected to immunohistochemistry targeting the residues T219 and Y221. These residues are known to be involved in the activation of ERK5 (Mody et al. 2003). Protein expression in epidermis and dermis was quantified separately. We could indeed observe enhanced staining of pERK5 in lesional skin compared to healthy skin in epidermis and dermis(Figure 2A,B).

**Figure 2.**
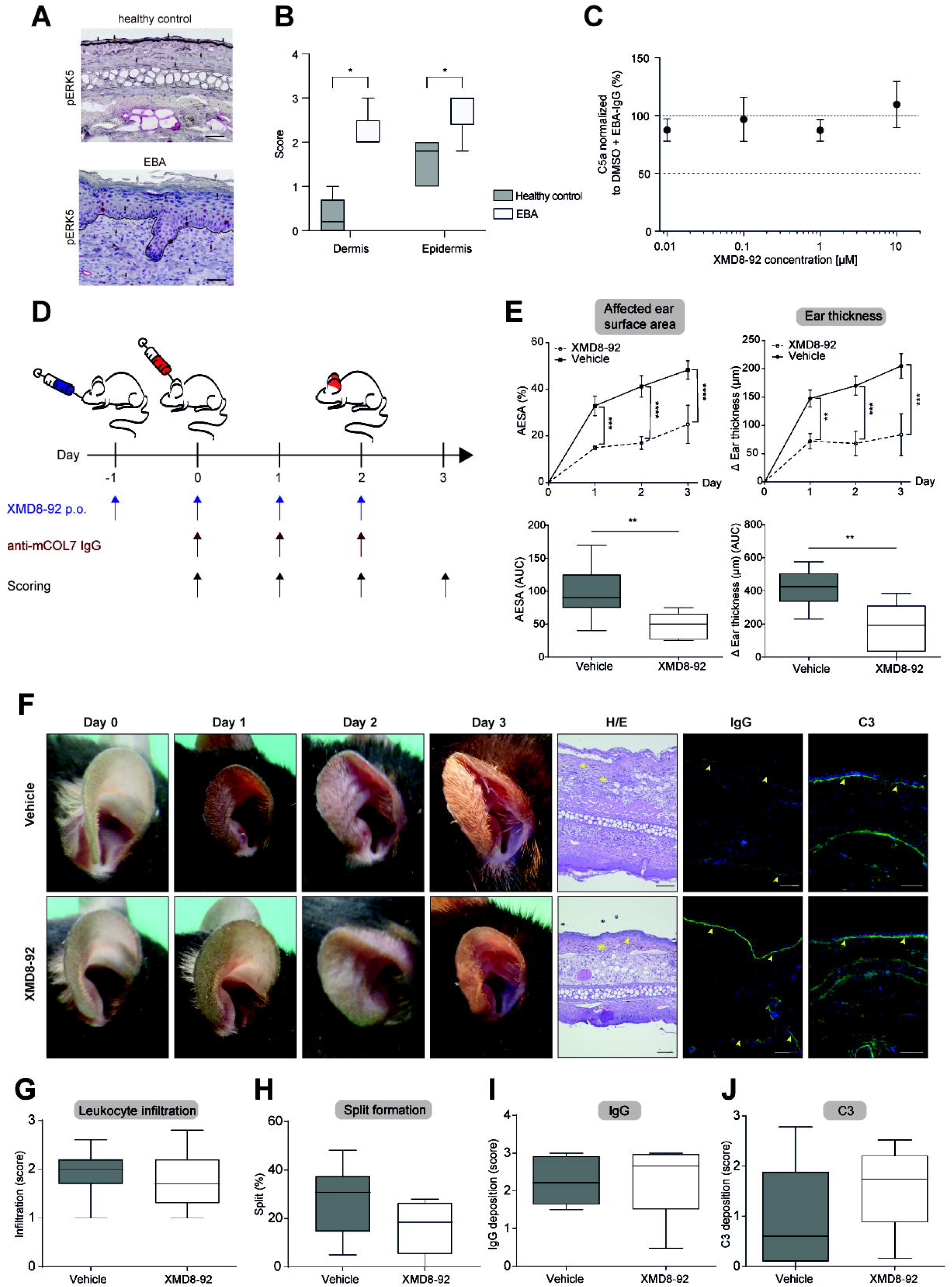
ERK5 is activated in lesional skin of Epidermolysis bullosa acquisita (EBA) *ex vivo*, and its inhibition by XMD8-92 alleviates disease development *in vivo* without affecting keratinocyte C5a release or antibody deposition *in vitro*. **(A)** Lesional and healthy skin obtained from immunization-induced EBA and healthy control mice was immunohistochemically stained for phosphorylated ERK5 (pERK5). Scale bar represents 100 µM. The dermal-epidermal junction is indicated by the black lines. Arrows indicate stained cells. **(B)** Enhanced staining in both dermis and epidermis in lesional treated mice was observed in a semi-quantitative measurement. **(C)** Immortalized keratinocytes (HaCaT) were stimulated with anti-COL7^C^ IgG and C5a concentration was measured using ELISA, which showed no significant changes upon ERK5 inhibition. **(D-F)** Antibody transfer–induced EBA was induced by injection of rabbit anti-mCOL7^C^ antibodies in C57BL/6J mice, and prophylactic treatment with XMD8-92 twice daily significantly reduced clinical disease scores, affected ear surface area (AESA), and delta ear thickness (referring to day 0) compared to vehicle control. **(G,H)** In skin samples obtained on the final day of the antibody transfer–induced EBA model, histological analyses revealed no significant effects of ERK5 inhibition on leukocyte infiltration (asterisk) or dermal–epidermal split formation (arrows). IgG or C3 deposition remained unchanged (arrows). **(I–J)** Cryosections were stained for IgG and C3 deposition, which were both unaffected by ERK5 inhibition. **B**: n = 5, Tukey boxplot Mann-Whitney test, *p ≤ 0.05; **C**: n = 3, , mean ± SEM ; **E**: n = 6 (control)/5 (XMD8-92), mean ± SEM, mixed effect analysis with Šidák’s multiple comparisons test, **p ≤ 0.01, ***p ≤0.001, ****p ≤ 0.0001; **G-J**: n = 6 (control)/5 (XMD8-92), Tukey boxplot, Mann-Whitney test

The presence of pERK5 in keratinocytes indicates the necessity of further investigation into the role of ERK5 in keratinocyte function since keratinocytes play a crucial role in attracting neutrophils to the skin by releasing inflammatory mediators such as C5a upon activation (Sadik et al. 2018; Sitaru et al. 2005). To evaluate the impact of ERK5 inhibition on C5a production, we again used the small-molecule inhibitor XMD8-92. The immortalized human keratinocyte cell line HaCaT was used to determine C5a levels by ELISA following stimulation with purified anti-COL7-IgG from EBA patients. No significant changes in C5a secretion were observed (Figure 2C). Therefore, C5a production by keratinocytes appears to be independent of ERK5.

Since the inhibition of ERK5 significantly reduced ROS release, a major hallmark of EBA pathogenesis (Chiriac et al. 2007b), we proceeded to evaluate XMD8-92’s inhibitory potential *in vivo*. XMD8-92 was administered in a prophylactic approach to assess its impact on disease development and progression of the antibody transfer-induced EBA model. Dose and treatment schedule were adapted from a previous publication using XMD8-92 (Yang et al. 2010). Briefly, C57BL/6J mice were injected with rabbit anti-mouse COL7^c^-IgG (Stüssel et al. 2020; Zillikens et al. 2021), and treated orally twice daily with XMD8-92 or vehicle, starting one day before disease induction, and subsequently, disease was scored daily (Figure 2D).

Clinical disease score and increase in ear thickness were significantly reduced at all time points (Figure 2E, F). On day 3, affected ear surface area (AESA) measured48.3 % in the vehicle group compared to 25.0 % in the treatment group. Ear thickness amounted to 250 µm in the vehicle group, compared to 83 µm in the treatment group. Histological examination revealed that leukocyte infiltration was unaffected by ERK5 inhibition (Figure 3G). Split formation showed a tendency to be reduced, however, this did not reach statistical significance (p = 0.12) (Figure 2 H). As anticipated, IgG and C3 deposition showed no significant differences between the groups (Figure 2I, J).

**Figure 3.**
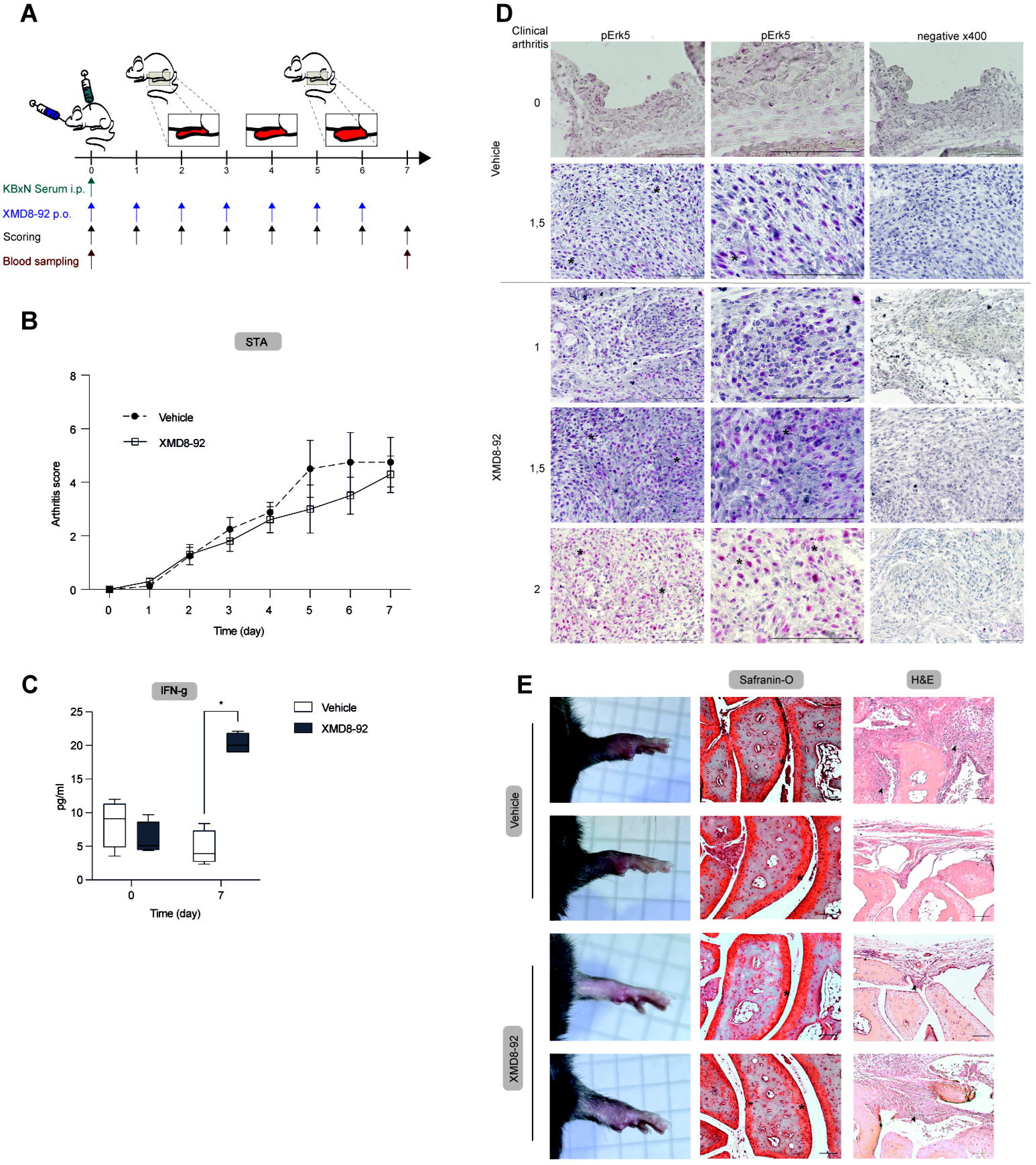
ERK5 inhibition does not affect serum-transfer arthritis(STA) severity. **(A)** Schematic overview of STA, which was induced in female C57BL/6J mice by injection of arthritogenic KBxN serum. Mice were treated with XMD8-92 twice daily. **(B)** Clinical arthritis scores were comparable between vehicle- and XMD8-92–treated groups over 7 days. **(C)** Serum samples were collected at day 0 and 7 and cytokine analysis revealed significant higher IFN-γ concentrations in XMD8-92–treated mice on day 7. Paws were collected on the final day of the experiment and **(D)** Immunohistochemistry demonstrated dense infiltrates of pERK5-positive cells in affected joints (asterisk) **(E)** Histological staining (H&E for leukocyte infiltration (arrows); Safranin-O for cartilage damage (asterisk)) reflected clinical disease scores. **B**: n = 4 (control)/5 (XMD8-92), mean ± SEM, n.s., mixed effect analysis with Šidák’s multiple comparisons test; **C**: n = 4, Tukey box plot, 2-way ANOVA, *p ≤ 0.05

### ERK5 inhibition does not alter STA disease severity

Given that the development of EBA, a neutrophil-dependent disease, was affected by ERK5 inhibition, we hypothesized that targeting ERK5 could similarly prove effective in other neutrophil-driven autoimmune diseases, such as RA. A commonly used model for RA is STA (Christensen et al. 2016), which was employed here. In contrast to EBA, mast cells, monocytes, macrophages and osteoclasts are also thought to play a major role in STA pathogenesis (Seeling et al. 2013). By an immunohistological staining of pERK5, we could demonstrate an increased expression of pERK5 in inflammatory infiltrates found in the legs of STA mice compared to healthy controls. (Figure 3A). Thus, we investigated whether ERK5 inhibition by XMD8-92 could improve the pathological phenotype similar to EBA. Therefore, STA was induced in C57BL/6J mice by injection of serum derived from arthritic KBxN mice. Joint swelling was monitored for 7 days and blood was sampled at day 0 and 7 to assess cytokine levels as an indicator of systemic inflammation (Figure 3B). No significant difference was observed between the two groups regarding the arthritis score (Figure 3C) but a significantly higher serum concentration of IFN-γ was identified, showing a four-fold increase in the treatment group compared to the control group after 7 days (Figure 3D). However, several other cytokines remained unchanged (Supplemental Figure 1). Histological stainings were performed to assess leukocyte infiltration (H&E) and cartilage damage (Safranin-O), and findings correlated with the varying clinical disease scores within treatment and vehicle group (Figure 3E).

### ERK5 inhibition does not affect platelet depletion in passive ITP model

To complement the *in vivo* analyses and gain insight into neutrophil specificity of ERK5 inhibition, XMD8-92 was tested in a non-neutrophil-dependent autoimmune disease model, specifically ITP. Similar to RA/STA and EBA, ITP is an autoantibody-mediated disease that can be accurately modelled experimentally, however, different cell types such as macrophages are involved in the effector phase. Platelet depletion was induced in C57BL/6J mice by i.p. injection of platelet-specific IgG2c. Platelet counts were measured prior to depletion and at 4 h and 24 h post-injection (Figure 4A). As expected, ERK5 inhibition did neither affect platelet depletion nor recovery (Figure 4B).

**Figure 4.**
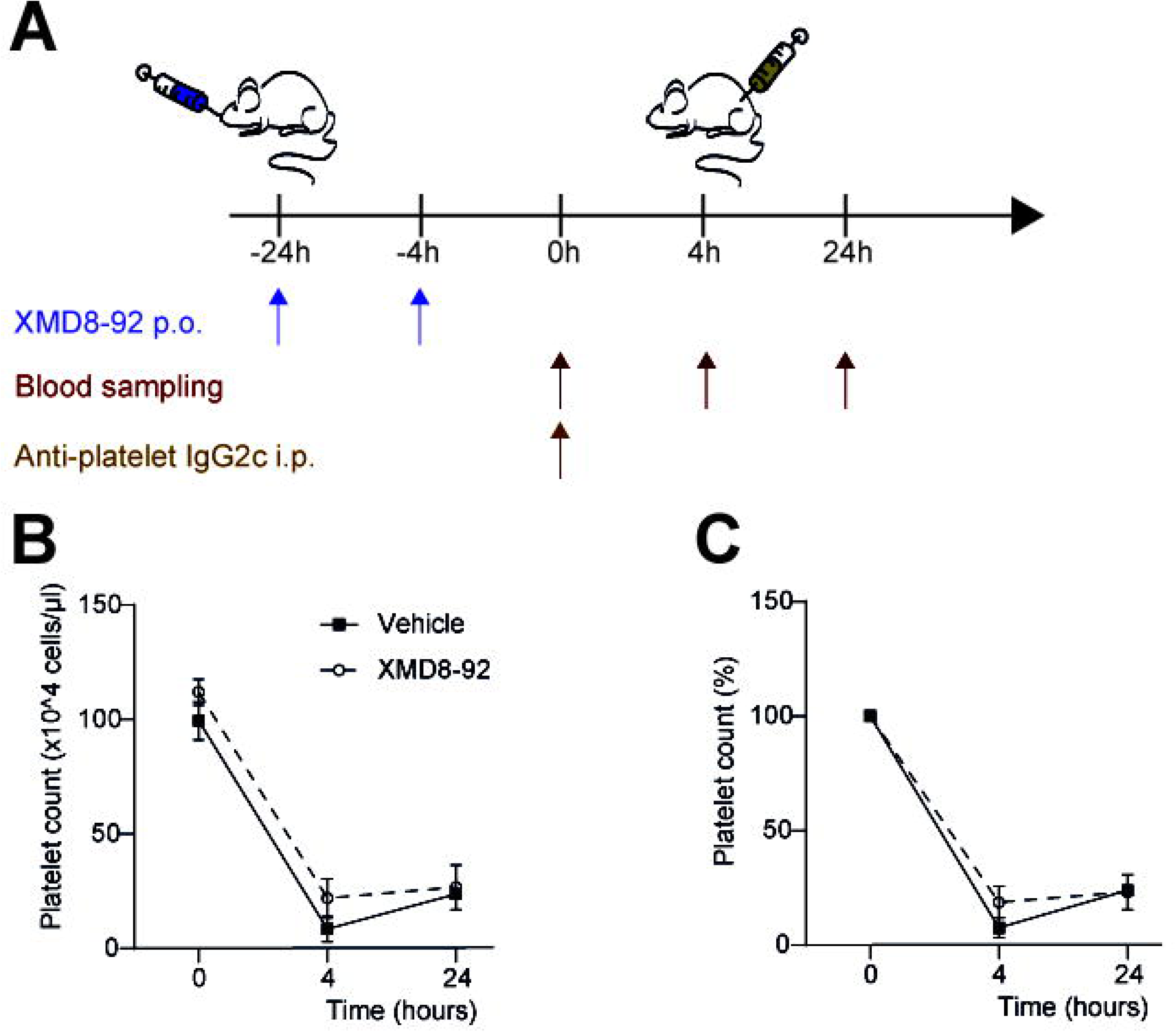
Treatment with XMD8-92 does not influence platelet depletion or recovery in immune thrombocytopenia (ITP). **(A)** ITP was induced in female C57BL/6J mice by i.p injection of anti-platelet IgG2c and platelet counts were measured at baseline and 4 and 24 h post-injection. **(B)** Absolute platelet count and **(C)** relative platelet count to baseline was comparable between XMD8-92- and vehicle-treated mice. **B, C**: n = 5 (control)/4 (XMD8-92), mean ± SEM, mixed effect analysis with Šidák’s multiple comparisons test.

## Discussion

Since current treatments for autoimmune diseases rely on general, untargeted immunosuppression with a low safety profile failing to induce remission in all patients (Lamberts et al. 2018), identifying additional therapeutic targets remains essential. In this study, we investigated the functional role of ERK5 in three prototypical autoantibody-mediated diseases, contrasting neutrophil-dependent and -independent ones. Neutrophils, as components of the innate immune system, function as key effector cells in many autoimmune diseases (Shimanovich et al. 2004). They cause tissue inflammation and damage by performing a variety of cellular functions, including adhesion, chemotaxis, cytokine release, NET formation, ROS production, and phagocytosis (Zhang et al. 2024). To broadly characterize neutrophils *in vitro*, we assessed, surface activation marker expression (CD18 and CD62L), adhesion, chemotaxis, and ROS release. ERK5 inhibition by XMD8-92 specifically reduced CD62L shedding and ROS release, without notably impacting viability. Although viability was significantly lower at the highest concentration tested, the percentage of viable cells was still in the range of the negative controls, and thus probably not of biological relevance. Likewise, CD18 expression, chemotaxis and adhesion were not affected, suggesting that later steps in neutrophil activation are affected. The ability of substances to inhibit ROS release *in vitro* indicates their potential to ameliorate ROS-dependent diseases *in vivo*, as demonstrated in both, the immunization-induced and the antibody transfer-induced EBA mouse models in recent studies (Ghorbanalipoor et al. 2024; Samavedam et al. 2018; Stüssel et al. 2020; Zillikens et al. 2021).

Activation of the ERK5 signaling pathway during neutrophil-dependent inflammation is *suggested* by the increased expression of pERK5, as observed in EBA-affected skin. In contrast, pharmacological ERK5 inhibition did not influence C5a secretion by HaCaT cells upon stimulation with anti-COL7-IgG.

In the antibody transfer-induced EBA mouse model, the systemic application of ERK5 inhibitor XMD8-92 alleviated disease progression from day 1 onwards. The degree of disease improvement was comparable to that achieved with treatments such as propranolol (Stüssel et al. 2020). As expected, IgG and complement component C3 deposition remained unchanged, indicating that the effect observed with ERK5 inhibition acts specifically during the effector phase. Furthermore, leukocyte infiltration was unaltered which is in line with the unchanged chemotaxis observed *in vitro.* Split formation, primarily driven by ROS release, showed a tendency to be reduced; however, this difference did not reach statistical significance.

To assess the impact of ERK5 inhibition across antibody-mediated neutrophil-dependent and -independent diseases, two additional murine models were included alongside EBA: STA and ITP. In the EBA model, as in KBxN-STA, neutrophils are the main cell type found in inflamed tissue. ITP on the other hand is completely independent of neutrophils, with macrophages being the main driver of pathogenesis (Krémer et al. 2022; Norris et al. 2023). As only antibody-transfer-induced EBA was ameliorated, it is important to emphasize the distinct pathogenic mechanisms underlying these disease models. Accordingly, the observed disease-specific efficacy of XMD8-92 suggests that the inhibitor indeed preferentially targets neutrophils and not cells of the mononuclear-phagocytic network. Alternatively, a major difference between passive EBA and STA to be considered is the activation of distinct signaling pathways, possibly leading to different effector mechanisms. In addition to neutrophils, macrophages and osteoclasts also contribute to STA pathogenesis. Furthermore, FcγRIII and FcγRIV signalling mediates inflammation in STA (Christensen et al. 2016), while tissue damage in EBA is mainly mediated via FcγRIV(Kasperkiewicz et al. 2012).

The hallmark of EBA is the release of reactive oxygen species (ROS). Here, neutrophil cytosolic factor 1 (Ncf1)-deficient mice, which cannot generate ROS through NADPH oxidase, are completely protected from disease manifestation (Chiriac et al. 2007b). Opposingly, as defects in nitrogen or ROS production could not rescue mice from developing arthritis, it has been hypothesized that NET formation and cytokine release might be more important for pathogenesis than ROS production (Wipke and Allen 2001). Moreover, Ncf1-deficient mice showed increased arthritis severity, which was partly attributed to the accumulation of neutrophils in the joints, along with upregulated pro-inflammatory characteristics (Liao et al. 2021). Interestingly, the source of ROS appears to be important, as macrophage-derived ROS can suppress T cell activation in an Ncf-1 dependent manner (Gelderman et al. 2007).

With cytokine production being a relevant factor for STA pathogenesis, systemic cytokine levels were measured before disease onset and at the peak of arthritis development. Albeit minor increases in IL-6 and IL-27 could be observed over the course of the disease, low overall concentrations of inflammatory cytokines in serum of STA mice are in line with the model primarily involving joint inflammation. Nevertheless, XMD8-92 treatment was associated with increased serum IFN-γ levels. Produced by Th1, CD8^+^ T cells and NK cells, IFN-γ appears to exhibit immunomodulatory functions in RA patients treated with anti-TNF treatment adalimumab (Aravena et al. 2011). In murine models of antigen-induced arthritis, IFN-γ deficiency has accordingly been associated with more severe disease (Irmler et al. 2007). While this supports a potentially disease-modifying role of XMD8-92 in arthritis, no significant clinical differences in disease progression were observed herein. The reasons behind this remain speculative and require further research.

In ITP on the other hand, interventions targeting the production of platelets, specifically megakaryocytes, or the effector cells in the reticuloendothelial organs (Krémer et al. 2022; Norris et al. 2023), would most likely lead to a reduction of the disease severity. Our data show that inhibition of ERK5 is negligible for this process. Interestingly, platelets themselves also express ERK5 (Cameron et al. 2015). Further investigation is necessitated, as other members of the ERK family have been implicated in regulating megakaryocyte differentiation (Mazharian et al. 2009; Séverin et al. 2010).

A limitation of our study is the exclusive use of an inhibitory compound rather than a targeted deletion of *Erk5*. Furthermore, XMD8-92 functions as a dual inhibitor, targeting both bromodomains (BRDs) and ERK5 (Lin et al. 2016). BRDs are conserved structural modules found in chromatin- and transcription-associated proteins, where they play a role in epigenetic regulation and transcriptional regulation by recognizing acetylated histones (Cochran et al. 2019; Mann et al. 2023; Wang et al. 2021). The effects of XMD8-92 may, in part, be due to its ability to inhibit BRD4. Therefore, further studies should be conducted using more selective ERK5 inhibitors to better isolate its specific effects (Lin et al. 2016). Studying ERK5’s isolated effects is challenging, as its substrates, the MEF2 transcription factors, can regulate myeloid cell fate by regulating the balance between monopoiesis and granulopoiesis (Schüler et al. 2008). Furthermore, we focused specifically on neutrophils but cannot exclude effects on other ERK5-expressing cells.

In our study, functional analysis of neutrophils revealed a specific role for ERK5 in ROS release, while other neutrophil functions following IC stimulation remained unaffected. Supporting the existing notion that the capacity to inhibit ROS release predicts *in vivo* efficacy, inhibition of ERK5 by XMD8-92 significantly reduced disease severity in the antibody transfer-induced EBA model. Contrarily, in STA and ITP ERK5 appears dispensable. These findings suggest that ERK5 plays a critical role in EBA pathogenesis, particularly in the effector phase by modulating ROS release from neutrophils.

## Materials and Methods

### Animal experimentation

C57BL/6J mice (Charles River, Germany) were bred in a specific pathogen–free environment and provided standard mouse chow and acidified drinking water *ad libitum*. Mice of both sexes, aged 8 to 14 weeks, were used for experimental models, with clinical examinations, biopsies, and bleeding procedures performed under anesthesia, either using isoflurane or intraperitoneal (i.p.) administration of a ketamine (75 mg/kg, Sigma-Aldrich, Taufkirchen, Germany) and medetomidine (1 mg/kg, Vetoquinol, Ismaning, Germany) mixture antagonised after ca. 30 min by atipamezole (5 mg/kg, Vetoquinol). On the final day, the mice were euthanized either using 15 mg/kg xylazine (Sigma-Aldrich) and 100 mg/kg ketamine and cervical dislocation or carbon dioxide.

### Sampling of human biomaterials

For the isolation of human neutrophils, whole blood was drawn from healthy volunteers after informed consent was given.

### Chemicals

All standard chemicals were supplied by Carl Roth (Karlsruhe, Germany) or Sigma-Aldrich (Taufkirchen, Germany). XMD8-92 was supplied by Selleckchem (Cologne, Germany) (Cambridge, UK). For *in vitro* assays, 10 mM XMD8-92 was dissolved in DMSO and further diluted to final concentrations of 1 µM, 0.1 µM, 0.01 µM and 10 µM. For *in vivo* experiments, XMD8-92 was dissolved in 30 % hydroxypropyl β-cyclodextrin, which also served as the vehicle control.

### Immunohistochemical staining of phosphorylated ERK5 (pERK5)

Paraffin-embedded skin specimens from control and immunization-induced EBA mice (Kasprick et al. 2019) or STA were used for immunohistochemical staining of pErk5. Immunohistochemistry was performed as previously described with modifications (Gross et al. 2024). Deparaffinization and rehydration were performed using standard protocols followed by heat-induced antigen retrieval in citrate buffer at pH 6 for 10 min at 110 °C. Samples were washed in tris-buffered saline (TBS) and permeabilized with 0.5 % (v/v) Triton X-100 in 6 % (w/v) bovine serum albumin (BSA) in TBS for 30 min at room temperature (RT). Blocking was performed with 10 % (v/v) normal goat serum in TBS for 30 min at 37 °C. Incubation with the primary antibody (anti-pErk5, PA5-114573, Invitrogen, Waltham, USA) diluted 1:200 in 2 % normal goat serum in TBS was performed for 1.5 h at RT. Subsequently, slides were incubated for 1 h at RT with goat anti-rabbit IgG H&L AP (ab97048, abcam) diluted 1:500 in 2 % normal goat serum in TBS. SIGMAFAST™ Fast Red (Sigma-Aldrich) served as substrate. Nuclei were counterstained with hematoxylin. Stained sections were evaluated by light microscopy (Keyence BZ-9000E, Osaka, Japan). Analysis was performed in a blinded, semi-quantitative manner, separately for the dermis and the epidermis. Expression was evaluated by visual inspection with scores of 0 (no staining), 1 (> 0 %, but ≤ 10 % positive cells), 2 (> 10 %, but ≤ 50 % positive cells), and 3 (> 50 % positive cells).

### Secretion of complement component C5a by HaCaTs

Immortalized keratinocyte (HaCaT) cells (Boukamp et al. 1988) were cultured with the cell culture medium Keratinocyte Growth Medium 2 (KGM2; Bio&Sell, Feucht, Germany) supplemented with Supplement Mix (PromoCell GmbH, Heidelberg, Germany), 0.06 mM CaCl_2_ (PromoCell GmbH, Heidelberg, Germany) and grown to confluency on 48-well plates at 5 × 10^4^ cells/well. After culturing for 18 h, HaCaT cells were pre-incubated with XMD8-92 at concentrations of 10 µM, 1 µM, 100 nM, 10 nM, or 1 nM (in 0.1 % DMSO) for 5 min followed by stimulation with affinity purified anti-COL7-IgG from EBA patients (80 µg/mL), as previously described.(Vorobyev et al. 2015; Zillikens et al. 2021) Cells stimulated with anti-COL7-IgG in absence of XMD8-92 (with 0.1 % DMSO) served as a positive control. Cells stimulated with human IVIG (Intratect, Biotest Pharma GmbH, Dreieich, Germany) in the presence of 10 µM of XMD8-92 served as negative control. Cells were incubated for 24 h, after which supernatants were removed and stored at −20 °C. C5a concentration in supernatants was determined using the Human Complement Component C5a DuoSet ELISA (R&D Systems GmbH, Wiesbaden, Germany) in accordance with manufacturer instructions and measured in a GloMax® Discover microplate reader (Promega, Walldorf, Germany).

### Human PMN purification

For ROS release assay, flow cytometry, and adhesion assay, PMNs were isolated from EDTA-treated human whole blood performing a PolymorphPrep™ (Serumwerk Bernburg AG, Bernburg, Germany) density gradient centrifugation as previously described (Stüssel et al. 2020). For the chemotaxis assay, PMNs were purified from citrated human whole blood using a Ficoll gradient (Skoog and Beck 1956). Blood was diluted 1:2 with 1 % (w/v) polyvinyl alcohol and allowed to sediment for 25 min. The supernatant was cautiously added to Ficoll-Paque™ PLUS (Cytiva, Marlborough, USA) and centrifuged for 24 min at 850 g without brake. Erythrocyte lysis was performed on the pellet using distilled water and stopped with 2x phosphate-buffered saline (PBS). After washing, the cells were diluted to 4 × 10^5^ cells/mL in 1 % (w/v) BSA in PBS. Neutrophil purity was assessed in flow cytometry for all experiments and was higher than 85 % for all samples.

### ROS release assay

Plates were coated with ICs consisting of human COL7 E-F at 2.5 µg/mL and anti-human COL7 IgG1 at 1.8 µg/mL as previously described (Recke et al. 2014; Yu et al. 2010). PMNs (2 × 10^5^ cells/well) were added in the presence or absence of XMD8-92. Only antibody (with cells), only antigen (with cells), or only cells served as negative controls, while PMA (10 nM/mL) served as a positive control. DMSO was adjusted to a final concentration of 0.01 %. All samples were measured in duplicates. Luminol was added to each well to reach a final concentration of 90 µg/mL and luminescence was measured every 2 min for 2 h at 37 °C using a GloMax® Discover microplate reader (Promega, Walldorf, Germany). The area under the curve (AUC) was calculated and values were normalized to IC-stimulated cells without XMD8-92.

### Adhesion assay

PMN adhesion was assessed using impedance-based real-time cell analysis as described previously (Yu et al. 2018). Briefly, a 96-well E-plate (Agilent, Santa Clara, USA) was coated with ICs as described for the ROS release assay. Inhibitor solutions were prepared and added to the plate before transferring PMNs (2 × 10^5^ cells/well). Impedance was measured every 2 min for 120 min using in an xCELLigence system (Agilent, Santa Clara, USA) and expressed as the cell index in arbitrary units. Data were analyzed as described for the ROS release assay.

### Chemotaxis assay

The chemotactic capacity of neutrophils was assessed using 48-well Boyden chambers (NeuroProbe Inc., Gaithersburg, USA) as described before(Boyden 1962; Stüssel et al. 2020). Human PMNs were isolated as previously stated. The lower chamber was blocked with 1 % (w/v) BSA in 0.1 M Na_2_CO_3_/0.1 M NaHCO_3_ (pH 9) for 1 h at 37 °C in a humid chamber. After buffer removal, 5 nM IL-8 (Peprotech, Hamburg, Germany) was loaded to the lower chamber and covered with a polycarbonate membrane (pore size 3 µm; Costar Nucleopore GmbH, Tübingen, Germany). The fully assembled chamber was incubated for 30 min at 37 °C. The PMN suspension was brought to a final concentration of 0.9 mM CaCl_2_ and 0.5 mM MgCl_2_ and pre-warmed in a water bath at 37 °C. XMD8-92 was added at the respective concentrations. The cell-inhibitor suspension was added to the upper chamber. Following incubation (1 h at 37 °C), myeloperoxidase assay was performed on the migrated cells. For this, the cells from the lower chamber were transferred to a microplate and lysed with an equal volume of 0.2 % hexadecyltrimethylammonium bromide. In parallel, a standard of lysed cells was applied to determine the cell number. TMB substrate (Promega) was added to the wells for approximately 2 min before stopping the reaction with 1 M H_2_SO_4_. Absorbance was read at 450 nm in a GloMax® Discover microplate reader (Promega, Walldorf, Germany).

### Flow cytometry

Human PMNs were isolated as previously described and stimulated with ICs, prepared in the same manner as for the ROS release assay, for 2 h at 37 °C in a 5 % CO□ atmosphere. Subsequently, they were stained for the flow cytometric analysis of the following surface markers using standard flow cytometry procedures: CD14 (clone HCD14), CD16 (clone 3G8), CD18 (clone 1B4/CD18), CD45 (clone HI30), and CD62L (clone DREG-56), CD193 (clone 5E8). Viability was assessed by Annexin V (B266195) and Zombie NIR staining (B308933).(Stüssel et al. 2020) All dyes were purchased from BioLegend (San Diego, USA). Measurements were performed on a MACSQuant® Analyzer 10 (Miltenyi Biotec, Bergisch Gladbach, Germany) and analysis was conducted using MACS Quantify (version 2.13.3, Miltenyi Biotec).

### Local antibody-transfer induced experimental EBA

Specific anti–mouse COL7^C^ IgG from rabbit serum was isolated as described previously (Bieber et al. 2017; Bieber et al. 2016; Sitaru et al. 2005). XMD8-92 was administered p.o. twice daily at 50 mg/kg body weight, beginning one day before the anti–mouse COL7^C^ IgG injection and was performed daily throughout the 5-day experiment. C57BL/6J mice of both sexes, aged 11-14 weeks, were injected in the base of the ear once with 100 μg rabbit anti–mouse COL7^C^ IgG. Mice were evaluated clinically every day, and the percentage of the affected ear surface area and ear thickness were assessed using a Mitutoyo 7301 dial thickness gauge (Neuss, Germany) by person unaware of the treatment. Ears were collected on the final day of the local antibody-transfer induced experimental EBA, cut in half and embedded in 4 % PBS-buffered paraffin or frozen in liquid nitrogen. Paraffin-embedded samples were subjected to hematoxylin-eosin staining. Split formation, cell infiltration, and epidermal thickness were evaluated as described before (Gross et al. 2024). Frozen samples were used for direct immunofluorescence staining detecting IgG and C3 deposits as described previously (Bieber et al. 2016).

### Serum-transfer arthritis (STA)

Serum transfer arthritis (STA) was induced by injection of arthritogenic KBxN serum in 8 week old female C57BL/6J mice at 10 µL/g bodyweight as described before (Seeling et al. 2023a). We limited analysis to female mice to reduce biological variability. Before and after arthritis induction, mice were treated twice daily with 50 mg/kg body weight XMD8-92 or vehicle control p.o. Severity of joint swelling was evaluated over the course of seven days by scoring the swelling of individual paws on a scale of 0-3. Scores were added for a total clinical score. Peripheral blood was collected retro-orbitally on day 0 and day 7 and inflammatory cytokines were quantified in the serum by LegendPlex multiplex assay (Mouse Inflammation 13-plex, BioLegend) as a parameter of systemic inflammation. On day 7, mice were sacrificed, and forelegs were prepared for histological analysis. Hematoxylin & eosin staining was performed to evaluate inflammatory infiltrates and Safranin-O staining to assess bone and cartilage damage, respectively, as previously described (Seeling et al. 2023b; Seeling et al. 2013). At the end of the *in vivo* arthritis experiments, both front paws of the mice were prepared and skin and surrounding muscle tissue were roughly removed. The paws were incubated overnight in 4 % PFA at 4 °C for fixation. Subsequently, bones were decalcified with Immunocal™ (Immunotec, Los Angeles, USA) for six days. For Safranin-O staining, nuclei were stained with Weigert solution (Weigert’s hematoxylin solution A and B 1:2, Carl Roth) for 3 min and washed in running water for 15 min. Sections were incubated in Scott’s buffer (1 % MgSO_4_, 0.2 % NaHCO_3_ in ddH_2_O), followed by water for 2 min each. Green color was achieved by incubation in Goldner solution III (Carl Roth), immersion in 1 % acetic acid and incubation in water for 30 s. Proteoglycans were stained with 0.1 % Safranin-O (Carl Roth) for 5 min. Sections were immersed in water, incubated for 1 min in 100 % ethanol and 3 min in xylol. Stained sections were evaluated by light microscopy using a Zeiss Axiovert 200 microscope and the AxioVision LE4.6 software (Olympus Life Science, Waltham, USA).

### Immune thrombocytopenia (ITP)

Platelet depletion in presence of kinase inhibition was assessed in 8 week old female C57BL/6J mice, as previously described (Wöhner et al. 2024). We limited analysis to female mice to reduce biological variability. Platelet depletion was induced by injection of 0.35 µg/g anti-platelet IgG2c (clone 6A6) upon treatment with vehicle control or XMD8-92 (50 mg/kg body weight applied p.o.) 24 h and 4 h before the induction of platelet depletion. Platelet counts in peripheral blood were assessed before 6A6-IgG2c injection, and at 4 h and 24 h post-injection in peripheral blood collected retro-orbitally. Platelet counts were determined using an ADVIA® 2120i hematocytometer (Siemens Healthineers, Germany).

### Statistical analysis

Data were analyzed using Prism 10.4.1 (GraphPad Software, San Diego, USA). Statistical tests were performed as indicated. p ≤ 0.05 was considered significant. (*p□≤□0.05; **p□≤□0.01; ***p□≤□0.001; ****p□≤□0.0001).

## Ethics statement and study approval

Animal experiments were approved by local authorities of the Animal Care and Use Committee (*Government of Schleswig-Holstein* and government of Lower Franconia) and performed by certified personnel (AZ 108/08-15 and *55.2.2-2532-2-2085*) following the ARRIVE guidelines. All experiments using human samples were approved by the local ethics committee (AZ #20-463, University of Lübeck, Lübeck, Germany) and were performed in accordance with the Declaration of Helsinki.

## Data availability

The data that support the findings of this study are available from the corresponding author upon reasonable request.

The authors declare that no conflict of interest exists.

## Acknowledgments

We thank Claudia Kauderer, Astrid Fischer and Alexandra Wobig for excellent technical assistance.

## CReDit author contributions

Conceptualization: AL, KB

Methodology: GV, RV, FP, XY, AR, NE, AL, KB

Investigation: SD, MS, LV, SJ, MR, CO, NG, MK, AS

Formal Analysis: SD, MS, LV, SJ, MR, CO, NG, MK, AS

Resources: AL, KB

Writing – Original Draft Preparation: SD, MS, AL, KB

Writing – Review & Editing: All authors

Supervision: GV, RV, FP, XY, AR, NE, AL, KB

Funding Acquisition: AL, KB

---

**Supplemental Figure 1: Most cytokine concentrations in serum samples collected at day 0 and 7 of serum transfer arthritis (STA) are unaffected by treatment with XMD8-92**. STA was induced by i.p. injection of arthritic KBxN serum. On day 0 and 7, serum samples were taken from STA and control mice. Cytokine levels were measured using a LegendPlex^TM^ multiplex assay. n = 4, Tukey box plot, 2-way ANOVA.

