## Supplementary figures and images for "ERK5 is required for neutrophil-mediated ROS release and a key contributor in epidermolysis bullosa acquisita"

### Supplemental Figure 1

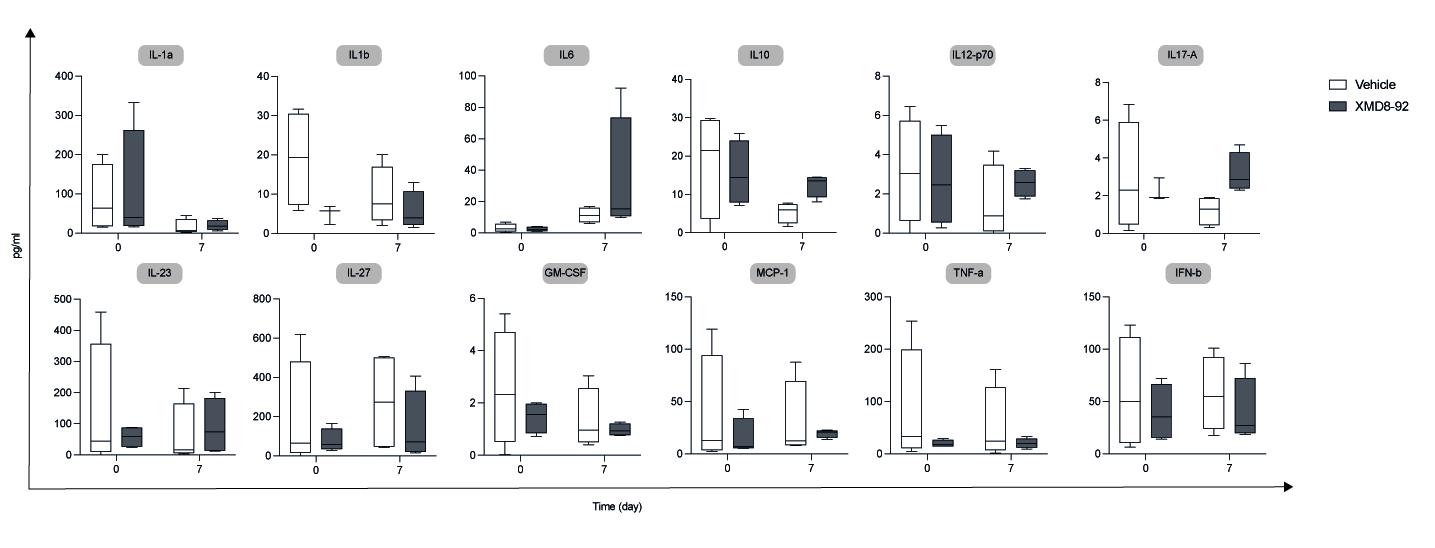
